# Validating methods to eradicate Select Agent and phylotype I *Ralstonia solanacearum* strains reveals that growth *in planta* increases bacterial stress tolerance

**DOI:** 10.1101/2022.03.15.484549

**Authors:** Madeline M. Hayes, Ronnie J. Dewberry, Lavanya Babujee, Rebecca Moritz, Caitilyn Allen

**Author notes:** Current address: Biosafety Director and Responsible Official, Colorado State University (Rebecca Moritz).

## Abstract

*Ralstonia solanacearum* is a destructive pathogen that causes bacterial wilt disease of diverse crops. Wilt disease prevention and management is difficult because *R. solanacearum* persists in soil, water, and plant material. Growers need practical methods to kill *R. solanacearum* in irrigation water, a common source of disease outbreaks. Additionally, the Race 3 biovar 2 (R3bv2) subgroup is a quarantine pest in many countries and a highly regulated U.S. Select Agent. Plant protection officials and researchers need validated protocols to eradicate *R. solanacearum* for regulatory compliance. To meet these needs, we measured survival of four R3bv2 and three phylotype I *R. solanacearum* strains following treatment with hydrogen peroxide, stabilized hydrogen peroxide (HuwaSan), active chlorine, heat, ultraviolet radiation, and desiccation. No surviving *R. solanacearum* cells were detected after cultured bacteria were exposed for ten minutes to 400 ppm hydrogen peroxide, 50 ppm HuwaSan, 50 ppm active chlorine, or temperatures above 50°C. *R. solanacearum* cells on agar plates were eradicated by 30s UV irradiation and killed by desiccation on most biotic and all abiotic surfaces tested. *R. solanacearum* did not survive the cell lysis steps of four nucleic acid extraction protocols. However, bacteria *in planta* were more difficult to kill. Stems of infected tomato plants contained a subpopulation of bacteria with increased tolerance of heat and UV light, but not oxidative stress. This result has significant management implications. We demonstrate the utility of these protocols for compliance with Select Agent research regulations and for management of a bacterial wilt outbreak in the field.

**Importance:** *Ralstonia solanacearum*, a globally distributed wilt pathogen of many high-value crops, is spread via diseased plant material and contaminated soil, tools, and irrigation water. The Race 3 biovar 2 Select Agent subgroup of *R. solanacearum* is subject to stringent and constantly evolving regulations intended to prevent pathogen introduction or release. We validated eradication and inactivation methods that can be used by: 1) growers seeking to disinfest water and manage bacterial wilt disease outbreaks; 2) researchers who must remain in compliance with regulations; and 3) regulators who are expected to define containment practices. Relevant to all these stakeholders, we show that while cultured *R. solanacearum* cells are sensitive to relatively low levels of oxidative chemicals, dessication, and heat, more aggressive treatment such as autoclaving or incineration is required to eradicate *R. solanacearum* cells growing inside plant material.

## Introduction

*Ralstonia solanacearum* is a globally distributed soil and water-borne plant pathogen. Collectively, this genetically diverse species complex can cause bacterial wilt disease in over 250 different plant hosts, including high-value agricultural exports and key subsistence crops like potato, tomato, and banana(1). *R. solanacearum* is transmitted via contaminated plant matter, soil, and water, making it very difficult to eradicate once it is established.

Though *R. solanacearum* is usually considered a tropical pathogen, a small subset of strains cause potato brown rot at temperatures as low as 20°C in tropical highlands and temperate zones(2). These strains fall into the phylotype II sequevar 1 subgroup, which is known historically and for regulatory purposes as Race 3 biovar 2 (R3B2)(3). Potato brown rot, which can cause up to 100% losses and is among the most destructive diseases of potatoes, can be disseminated via latently infected seed potato tubers and ornamental cuttings(4, 5). Regulatory agencies hope to prevent the spread of potato brown rot in higher latitudes, so *R. solanacearum* R3b2 is listed as a high-concern quarantine pest in Canada and Europe and is a Select Agent (SA) pathogen in the U.S. (6).

The Select Agent list includes microbes considered potential bioterrorism threats to humans, livestock, or crops. SA regulations evolve continuously to manage threats of pathogen release from accidental error, inadequate decontamination, or security flaws. For example, following an accidental release of live anthrax spores from an SA laboratory, SA regulations now require validated protocols that ensure all Select Agents are biologically inactivated before downstream applications such as experiments with extracted proteins or nucleic acids(7). Institutional SA programs throughout the US work closely with research labs to identify such applications and ensure that each inactivation protocol is appropriately validated.

Standardized inactivation protocols can prevent spread of pathogens, keep research labs in regulatory compliance, and help growers manage disease. However, a literature survey found no validated chemical and physical treatment methods to eliminate *R. solanacearum* from cultures, contaminated soil, water, or plant material.

To address this need, we defined treatments that will inactivate *R. solanacearum* in laboratory and agricultural conditions. We identified the levels of bleach (active Cl^−^), oxidative stress, ultraviolet (UV) radiation, heat, and desiccation required to eradicate *R. solanacearum* from water, plant matter, and laboratory surfaces. Treatments were validated on seven *R. solanacearum* strains, including four R3bv2 SA strains and three strains isolated from recent bacterial wilt outbreaks. Additionally, we validated that no viable *R. solanacearum* cells survived four protocols for extraction of nucleic acids, which are commonly used for downstream research and diagnostic purposes. There were no differences in treatment efficacy between SA and non-SA strains. However, a subpopulation of *R. solanacearum* cells growing in plants are highly resistant to heat and UV radiation. This finding has significant implication for *R. solanacearum* eradication protocols and effective bacterial wilt disease management.

## Materials and Methods

### Strain Selection

Strains used in these studies are described in Table 1. Four R3b2 *R. solanacearum* strains isolated from diverse geographical locations were chosen to represent the SA subgroup. Three additional *R. solanacearum* phylotype I strains were isolated from plant material from agricultural operations experiencing bacterial wilt outbreaks.

**Table 1:**
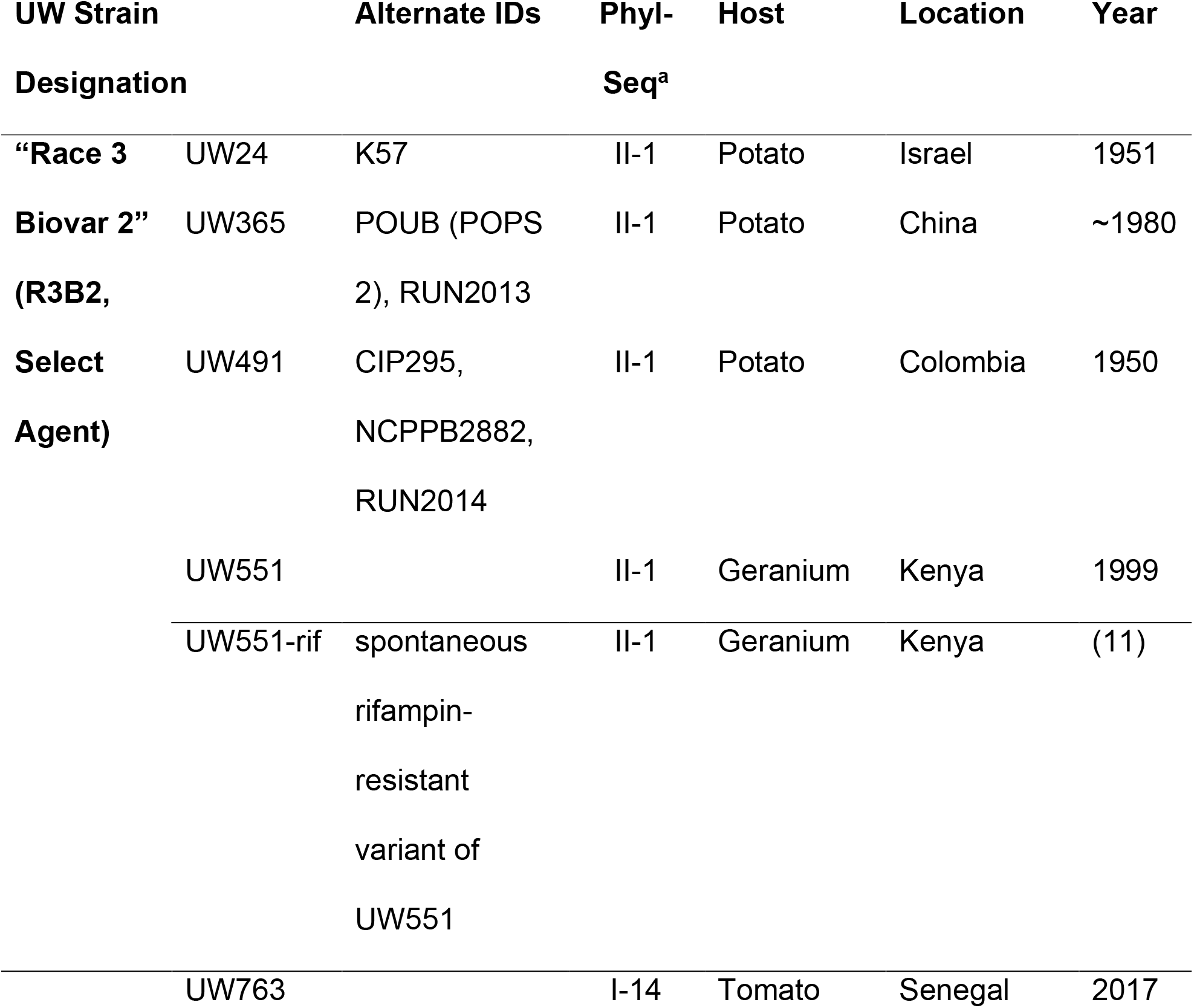
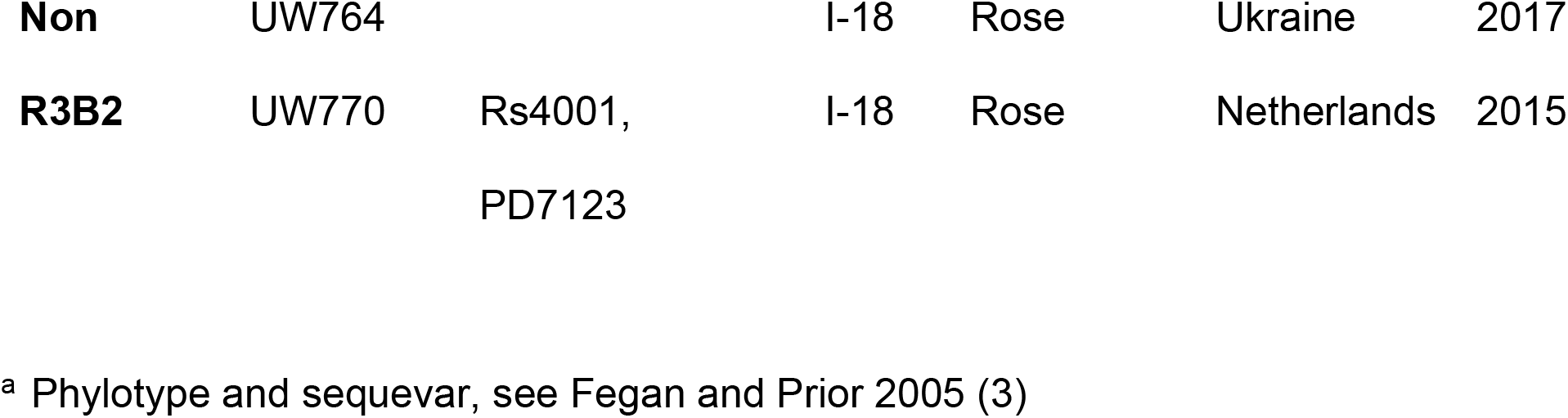
Characteristics of *R. solanacearum* strains used in this research.

### Preparation of Cultured Bacterial Cells for Treatment

Bacterial strains were cultured from water stocks on CPG+TZC agar plates (+ 50 mg/L rifampicin for UW551-rif) for 48 h at 28°C(8). Individual colonies were used to inoculate CPG broth, which was incubated overnight at 28°C with shaking at 225 rpm. After 24 h, cells were pelleted at 6800 x *g* RCF and resuspended in sterile water at OD_600_=0.8 (~1×10^10^ CFU/ml). This high-density bacterial cell suspension was then subjected to treatments as described below.

### Quantifying *R. solanacearum* Survival

#### Dilution plating

Treated cell suspensions were serially ten-fold diluted and plated in triplicate on CPG+TZC agar plates (+rifampicin for UW551-rif), incubated at 28°C for 48h, and colonies were counted for dilutions showing 3-30 colonies to determine the CFU/mL in the original suspension. The average of this triplicate plating was considered a single technical replicate. This method of dilution plating has a limit of detection of 100 CFU/mL.

#### Enrichment Culture

To determine if treated sample contained viable cells below the limit of detection for dilution plating, the remainder of each treated sample was enrichment cultured overnight at 28°C with 225 rpm shaking in 5 mL of CPG broth or, for soil or plant samples, in SMSA semi-selective media(9). If culture growth was visible, 500μL of this culture was centrifuged at 6000 rpm for 5 min in a 1.5 mL microcentrifuge tube. To determine if the growth contained *R. solanacearum*, the pellet was tested with the Agdia Rs immunostrip test (Agdia, Inc., Elkhart, IN, catalogue #ISK 33900) according to the manufacturer’s instructions. The limit of *R. solanacearum* detection following this enrichment culture method is 10 CFU/ml(10).

### Preparation of Bacterial Cells from Plants and Biofilms

Unwounded 21-day old tomato plants (bacterial wilt-susceptible cultivar ‘Bonny Best’), grown in 4-inch pots at 28°C with a 12h light/dark cycle, were inoculated using by soil soak(8). Briefly, 50 mL of a stationary phase bacterial culture, suspended in water as described above, was poured onto the soil of each pot and symptoms were allowed to develop. To assess behavior of *R. solanacearum* cells growing *in planta*, two 0.25 g stem sections were collected from tomato plants at disease index 1-2 (1-50% of leaves showing bacterial wilt symptoms), with the stem sampled between the petiole and first true leaf.

#### Planktonic cell extraction from infected plant tissue

Stem sections from infected plants were placed vertically in a 1.5-ml microcentrifuge tube containing 750 μL sterile H_2_O and centrifuged at 17.5x*g* for 10 min to remove unattached planktonic cells from the xylem vessels and leave behind cells in biofilms. The resulting cell pellet was vortexed to resuspend cells and the suspension was subjected to eradication treatments and bacterial survival was quantified as described above.

#### Growth from biofilms on glass slides

*R. solanacearum* was cultured as described above and suspended in liquid CPG media at OD_600_=0.1 (~ 1×10^7^ CFU/ml). This suspension was then used to inoculate glass well chamber slides as described previously (TRAN et. al.) to a final concentration of 1×10^6^ CFU/ml. Briefly, inoculated slides were incubated statically at 28°C for 72 h, with media decanted and replaced every 24 h. Slides were then dipped in sterile water to rinse away unattached cells. As a control to confirm the presence of a biofilm, a subset of rinsed slides were stained with crystal violet and evaluated microscopically. Experimental slides were placed in 50°C CPG media for 10 min, then immediately transferred to a room temperature bath to cool. These were then subjected to enrichment culture and bacterial survival was qualitatively assessed as described above. For UW763, three biological replicates with three technical replicates each were performed (total n=9). For UW551, one biological replicate with three technical replicates each was performed (total n=3). Strain UW551 formed little biofilm on glass slides under these conditions.

### Eradication Treatments

#### Chemical Treatments

Hydrogen peroxide (laboratory grade 30% vol/vol H_2_O_2_ solution, Avantor catalogue # 5240-02), HuwaSan stabilized peroxide (50% solution, Roam Technology, Ghent, Belgium), and commercial bleach (8.25% sodium hypochlorite, Clorox catalogue #30966) were added to a 5mL suspension of *R. solanacearum* cells to achieve the desired concentration, mixed thoroughly by vortexing, and allowed to stand at room temperature for 10 min. Cell survival was then quantified by serial dilution plating as described above. For each strain, a total of three biological replicates with three (H_2_O_2_ and HuwaSan) or one (bleach) technical replicates each (total n= 3 to 9) was performed.

#### Heat

100 μL of *R. solanacearum* cell suspension was transferred to 0.2-mL PCR tubes and incubated at various temperatures in a MJ Research PTC-200 gradient thermal cycler for various times. Cell survival was then quantified by serial dilution plating as described above. For each strain, three biological replicates were performed with one technical replicate each (total n=3).

#### In planta heat

Infected tomato stem samples obtained as described above were cut into 0.1 g stem sections, placed in a 2.0-mL homogenizer tube with 900 μL sterile water and four stainless steel beads and ground in a Qiagen PowerLyzer 24 at 2200 rpm for 1.5 min with 4 min rest for 2 cycles. Homogenate was then exposed to heat or room temperature and serially dilution plated as described above to determine CFU/g stem. For each strain, three biological replicates were performed with variable technical replicates (total n=5-45).

#### In planta desiccation

Infected tomato stem samples obtained as described above were desiccated in a commercial food dryer (Magic Mill MFD-9050) for 12 h at 35°C. *R. solanacearum* survival was assessed qualitatively by enrichment culturing the desiccated plant stem in 5 mL SMSA semi-selective media as described above. For each strain, two to three biological replicates were performed with variable technical replicates (total n=8-28).

#### In soil desiccation

Plants inoculated with UW551-rif were allowed to wilt and die in the pot, releasing *R. solanacearum* cells from the roots. A 1-2 g soil sample was removed from each pot and 0.1-0.2 g of this soil was cultured overnight at 28°C in SMSA+rif broth and qualitatively assessed for *R. solanacearum* growth with an Agdia Rs immonostrip to establish an initial +/− baseline for surviving *R. solanacearum*. The remaining soil sample was then dried as described above. Survival was determined qualitatively by enrichment culture of approximately 0.1-0.2 g soil in 5 mL SMSA semi-selective media as described below. Three biological replicates were performed with two technical replicates each. The spontaneous rifampicin resistant UW551(11) mutant, previously determined to have wildtype virulence and *in-planta* fitness, was used to increase selectivity from the dense soil microbiome.

#### Desiccation in tubes

0.5 mL of bacterial cell suspension was transferred to 1.5-mL polyethylene microcentrifuge tubes and allowed to stand (lid open) at 28°C for 120h. 0.5 mL sterile water was added to the tube and *R. solanacearum* survival was quantified by serial dilution plating as described below. For each strain, a total of three biological replicates were performed with one technical replicate each (total n=3).

#### *Desiccation on laboratory surfaces* (also known as fomites)

Common laboratory paper towels, chosen because they are used to contain spills, were tested for ability to harbor live cells after desiccation. 100 μL of cell suspension was placed on sterilized brown paper towel [Schilling Supply Company, catalogue #CS1751] and allowed to stand in a lidded but slightly open sterile petri dish at 28°C for 120 h. Fomite was then used to inoculate 5 mL liquid CPG media as described above in “Enrichment Culture.” For each strain, a total of three biological replicates were performed with three technical replicates each (total n=9).

#### Ultraviolet Light

*R. solanacearum* cell suspension was serially diluted onto solid CPG media plates and liquid was allowed to absorb completely into the agar. Uncovered plates were then exposed to 0.2 umol/m^2^/s-1 of radiation (254 nM wavelength) delivered by the germicidal UV lamp of a Baker SterilGARD®III Advance Biological Safety Cabinet (delivered UV radiation intensity was measured using a UV meter (Apogee Technologies MQ-200). For each strain, a total of three biological replicates were performed with one technical replicate each (total n=3).

### Select Agent Inactivation Treatments

We tested the ability of four nucleic acid extraction protocols to inactivate *R. solanacearum* as required for compliance with Select Agent regulations. Briefly, cells of R3bv2 strain UW551 suspended as described above were subjected to the inactivation (cell lysis) step of the following protocols: 1) Qiagen RNEasy RNA extraction kit (catalogue #74104): Pellet from 0.5 ml cell suspension was resuspended in in 450μL Buffer RLT according to the manufacturer’s instructions; 2) Epicentre MasterPure DNA extraction kit (catalogue #MC85200): Pellet from 0.5 ml cell suspension was resuspended in 400μL C&T Lysis buffer according to the manufacturer’s instructions; 3) Promega Maxwell DNA Extraction using the RSC Cultured Cells DNA cartridges (catalogue #AS1620) according to the manufacturer’s instructions, with cell viability determined 100 sec after bacteria were added to the cartridge; and, 4) Phenol-chloroform extraction method: Pellet from 0.5 ml cell suspension was resuspended in 5% water-saturated phenol/95% ethanol stop/lysis solution. Following these cell lysis steps, *R. solanacearum* survival was quantified as described above. Inactivation treatments were performed only on UW551, and a total of three biological replicates were performed with one technical replicate each (total n=3).

#### Experimental Design and Data Analysis

Each experiment contained multiple biological and technical replicates. For in culture assays, each biological replicate consisted of a single overnight culture which was inoculated with a single isolated colony from an agar plate. Technical replicates were repeats of each experiment using the same starting culture and suspension (i.e. one culture divided into three suspensions was one biological replicate with three technical replicates). For *in planta* experiments, one biological replicate consisted of all plants treated with inoculum from an overnight culture grown from a single colony as described above. One technical replicate was a single plant inoculated with that starting culture (i.e. one culture used to generate inoculum for fifteen plants was one biological replicate with fifteen technical replicates). Data were analyzed using Prism.

## Results

To determine the limits of survival for diverse *R. solanacearum* strains under field and laboratory conditions, we determined the lethal doses of a diverse set of chemical and physical treatments. LD_100_ was defined as the treatment necessary to reduce surviving cells below our 10 CFU/mL limit of detection. We tested treatments on 10^10^ CFU/mL cell suspensions because such dense cultures are common in the laboratory. This is much higher than the 10^2^ to 10^5^ CFU/g observed in naturally infested field soils or water, but any treatment effective at 10^10^ CFU/mL was also effective at lower densities (data not shown). *R. solanacearum* cells in infected plants were subjected to eradication treatments when wilt symptoms first appeared, as would happen in the field when growers try to control disease.

### Cultured *R. solanacearum* cells are sensitive to antimicrobial oxidants

We measured the ability of three antimicrobial chemicals, hydrogen peroxide, HuwaSan, and bleach, to kill cultured *R. solanacearum* cells in water. Ten minutes’ exposure to 400 ppm hydrogen peroxide reduced populations of all tested strains to below the limit of detection (Fig. 1A). Strains UW24, UW365, UW764, and UW770 did not survive 200 ppm hydrogen peroxide for 10 min. Treating bacteria with 50 ppm HuwaSan, a chemically stabilized form of hydrogen peroxide, also left no detectable CFU of any strain (Fig. 1B). Some strains were more sensitive to HuwaSan; 25 ppm eliminated UW491, UW763, and UW764 and only 12.5 ppm left no detectable CFU of strains UW365 and UW770. Active chlorine was highly effective against *R. solanacearum*. Ten minutes’ exposure to 0.1% v/v commercial bleach solution (8.25% sodium hypochlorite) in water, equivalent to 120 μM active chloride ion, left no detectable cells of all tested *R. solanacearum* strains (Fig. 1C). In summary, relatively low concentrations of these oxidative stress-inducing chemicals killed *R. solanacearum* cells. The LD_100_ of these antimicrobial chemicals was not affected by culture growth phase, with similar efficacy against log phase and stationary phase cells (data not shown).

**Figure 1.**
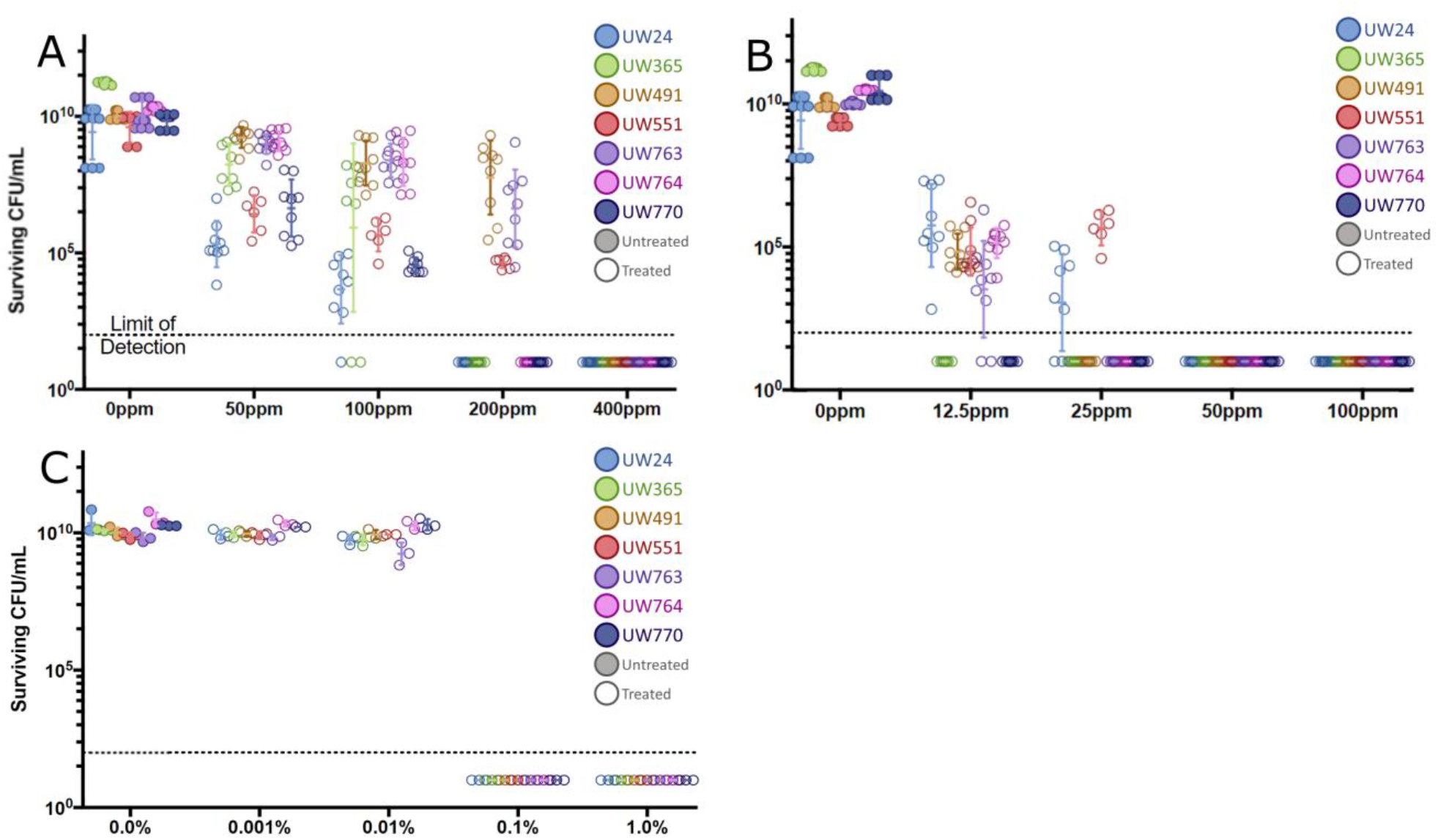
Oxidant concentrations required to eradicate *R. solanacearum*. Population sizes of seven *R. solanacearum* strains (identified by colored symbol as indicated in legend) following exposure to antimicrobial oxidants. Each symbol represents the result of one independent experiment, each containing three technical replicates. Surviving cells were quantified by serial dilution plating after a 10^10^ CFU/ml *R. solanacearum* cell suspension in water was incubated for 10 min at room temperature with indicated concentrations of **A. laboratory grade H_2_O_2_** (N = 9 experiments per strain); **B: 50% HuwaSan solution** (N = 9 experiments per strain); or **C: sodium hypochlorite**(N= 3 experiments per strain)

### Efficacy of physical treatments against *R. solanacearum*

Ultraviolet (UV) radiation was effective against cultured *R. solanacearum* cells spread on agar plates. Irradiation for 20 s reduced survival to below the limit of detection of all *R. solanacearum* strains except UW365, which required 30 s exposure (Fig. 2). This indicates that germicidal UV radiation produced by a standard biological safety cabinet can decontaminate *R. solanacearum*-infested surfaces.

**Figure 2:**
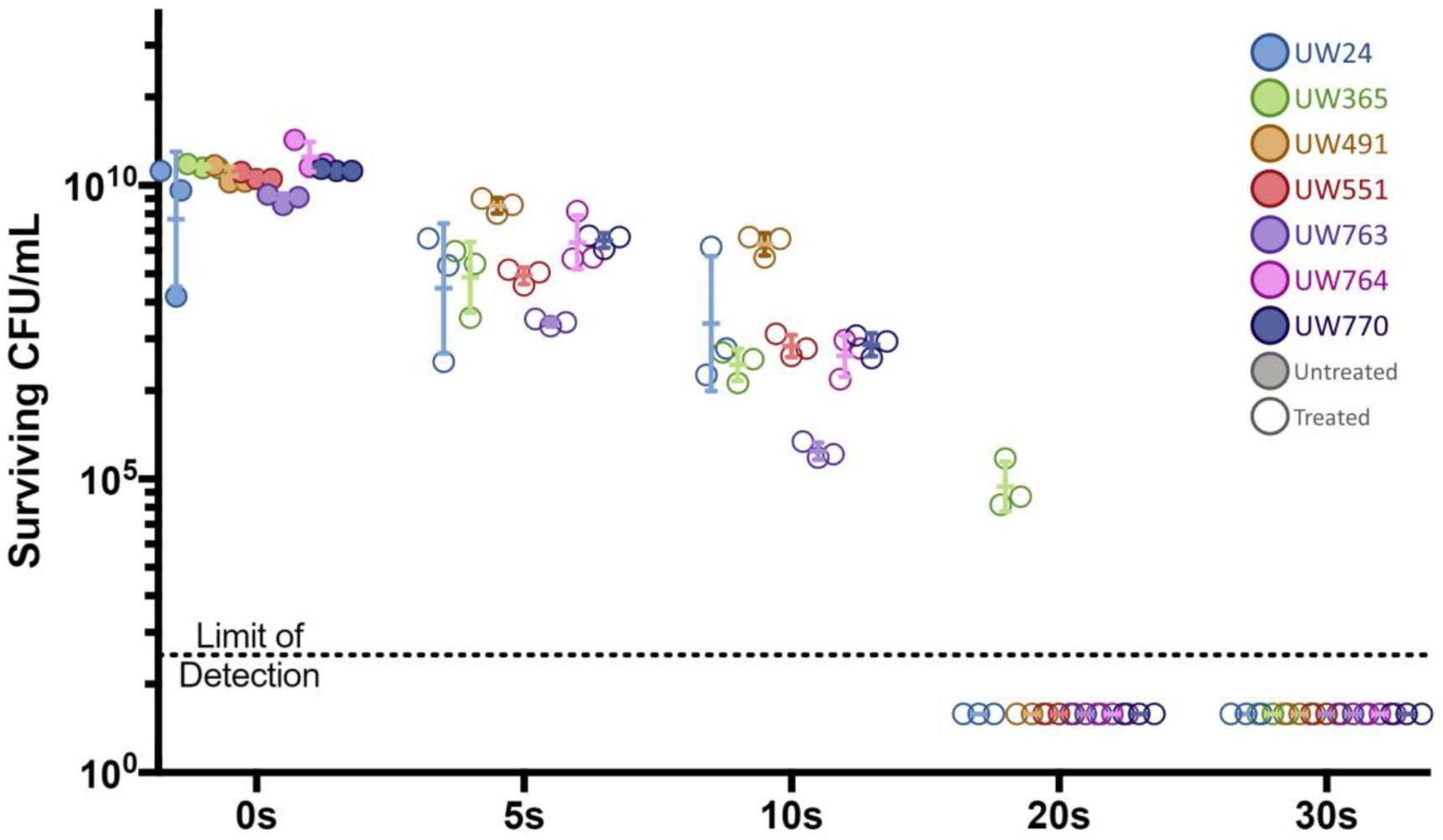
Ultraviolet radiation required to eradicate *R. solanacearum* from surfaces. Population sizes of seven *R. solanacearum* strains (identified by colored symbol as indicated in legend) following exposure to 0.2 umol/m^2^/sec UV light on CPG agar plates for the indicated time. Surviving cells were quantified by counting colonies that grew after treatment. Each symbol represents one independent experiment with three technical replicates (N = 3 experiments per strain) Desiccation is not usually a practical decontamination procedure, but it is useful to know a pathogen’s desiccation tolerance when cleaning growth facilities or following contaminated materials to disposal. We asked if a dry paper towel or other fomite (surface) can harbor viable pathogen cells and pose a risk for release. We measured bacterial survival after *R. solanacearum* cell suspensions were dried at 28°C for 120 h in open microcentrifuge tubes, then cultured in rich broth. None of the seven tested strains survived this treatment. Similarly, none of the strains survived 120 h desiccation on sterile laboratory paper towels, indicating that cultured *R. solanacearum* cells are intolerant of desiccation (Table 2).

Heat, usually autoclaving, is a common laboratory decontamination procedure. We measured heat tolerance of *R. solanacearum* cells grown under various conditions, as summarized in Fig 3. We determined that ten minutes’ exposure to 50°C is the minimum kill temperature for a water suspension of cultured *R. solanacearum* cells of any tested strain. This exact kill point was identified using a heat gradient that left surviving cells at 49.2°C but none at 51.4°C (Fig 4A). Thus, cultured *R. solanacearum* cells are efficiently killed by brief exposure to relatively low temperatures.

**Table 2:** Desiccation on surfaces and in plant tissue and soil eradicates *R. solanacearum*.

| Strain<sup>a</sup> | *In planta*<sup>b</sup> | In soil<sup>e</sup> | In tubes <sup>f</sup> | On fomite<sup>g</sup> |
| --- | --- | --- | --- | --- |
| UW24 | 0/14<sup>c</sup> (n=3) <sup>d</sup> | NT | 0/3 (n=3) | 0/9 (n=3) |
| UW365 | 0/13 (n=3) | NT | 0/3 (n=3) | 0/9 (n=3) |
| UW 491 | 0/8 (n=2) | NT | 0/3 (n=3) | 0/9 (n=3) |
| UW 551 | 1/28 (n=3) | 0/6 (n=3) | 0/3 (n=3) | 0/9 (n=3) |
| UW 763 | 1/19 (n=3) | NT | 0/3 (n=3) | 0/9 (n=3) |
| UW 764 | 0/17 (n=2) | NT | 0/3 (n=3) | 0/9 (n=3) |
| UW 770 | 0/22 (n=3) | NT | 0/3 (n=3) | 0/9 (n=3) |

<sup>a</sup> See Table 1 for details
<sup>b</sup> Infected plant stems were dehydrated at 35°C for 12 hours, then used to inoculate SMSA liquid media, enrichment cultured at 28°C for 48 hours, then qualitatively assessed for bacterial growth.
<sup>c</sup> Each cell shows number of technical replicates showing survival out of (/) total technical replicates tested
<sup>d</sup> n= number of biological replicates
<sup>e</sup> Soil from pots of completely wilted plants dehydrated and then assessed for survival as with infected plant stems described above
<sup>f</sup> Cell suspensions in water (aliquoted into 1.5mL tubes) allowed to desiccate at 28°C for 120 hours, then used to inoculate CPG liquid media and enrichment cultured at 28°C for 48 hours, then qualitatively assessed for growth
<sup>g</sup> Cell suspensions in water were dropped onto paper towels and allowed to desiccate then assessed for survival with enrichment culture as with suspensions in tubes described above

**Fig. 3.**
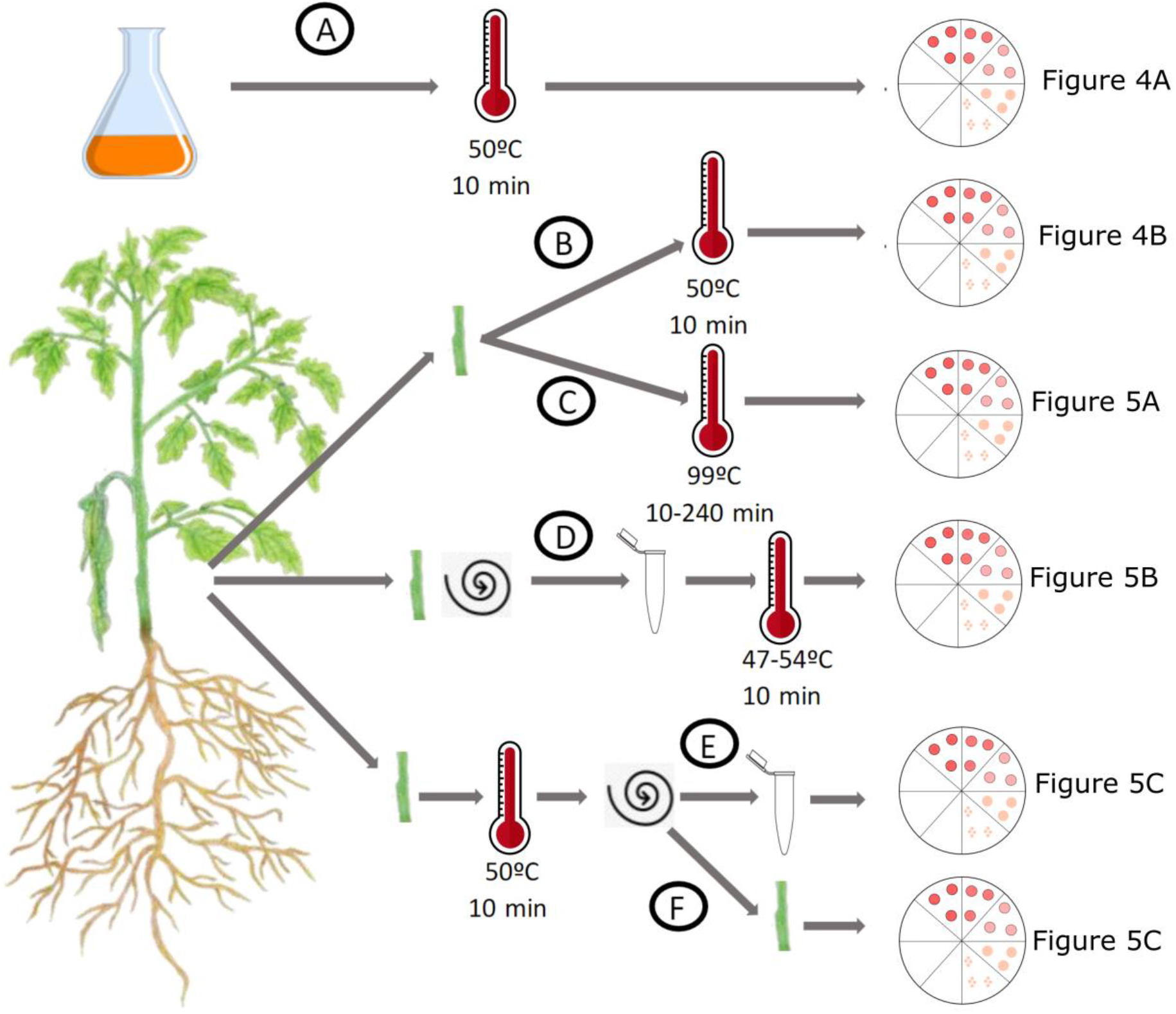
Overview of experiments to determine heat tolerance of *R. solanacearum* cells in culture and from infected plants. For all treatments, surviving cells were quantified by serial dilution plating combined with a qualitative enrichment culture to check for surviving cells below the 100 CFU detection limit of dilution plating. Quantitative results for each experiment are displayed in the figure referenced. **A**. Surviving *R. solanacearum* cells after 100 μL of a 10^8^ CFU/ml suspension of cultured cells in a 0.2 mL PCR tube was exposed to 50°C for 10 minutes. **B**. Surviving *R. solanacearum* cells after stem sections from infected tomato plants were exposed to 50°C for 10 minutes. **C**. Surviving *R. solanacearum* cells after stem sections from infected plants were exposed to 99°C for 10 minutes. **D**. Surviving cells after planktonic *R. solanacearum* cells were extracted from infected tomato stems by gentle centrifugation, then exposed to 47 to 54°C for 10min. **E-F)** Surviving *R. solanacearum* cells after stem sections from infected plant tissue were treated at 50°C for 10 minutes, then gently centrifuged to extract planktonic cells. Both the surviving extracted cells (E) and surviving cells in the ground plant stem (F) were quantified.

**Figure 4.**
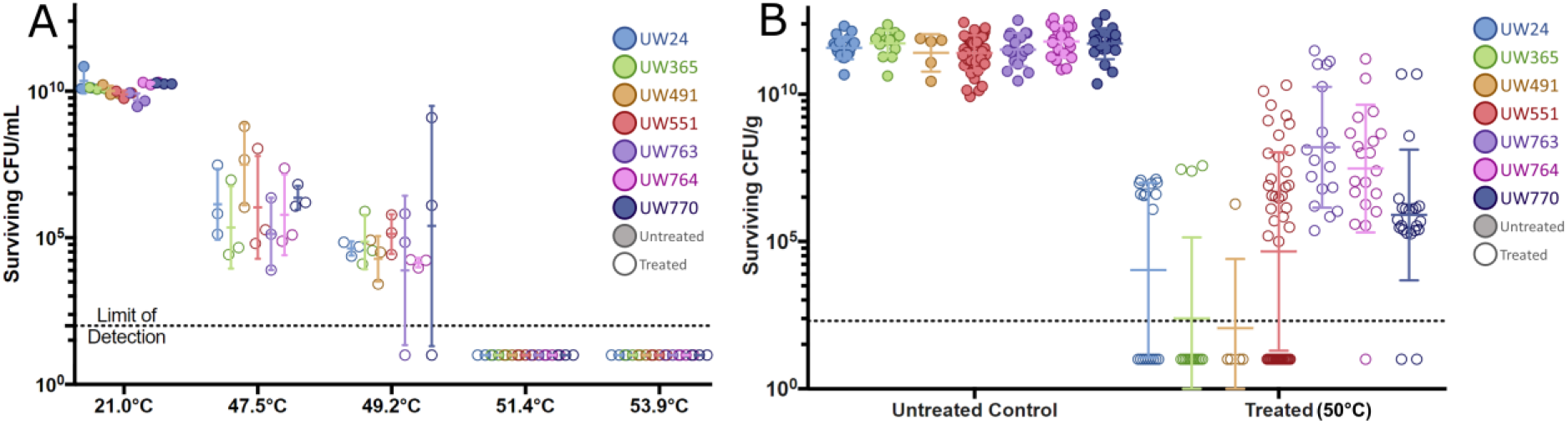
Heat treatments required to eradicate *R. solanacearum* cells from culture or from infected plants. **A. Cultured** *R. solanacearum* **cells are very susceptible to heat**. Population sizes of seven *R. solanacearum* strains (indicated by colored symbol) following exposure of suspensions of cultured cells in water at the indicated temperature for 10 min. Surviving cells were quantified by serial dilution plating. (N = 3 experiments per strain) **B**. *R. solanacearum* **cells in infected plant tissue are more resilient to heat stress**Population sizes of seven strains (indicated by colored symbol) following treatment of stems from *R. solanacearum*-infected tomato plants at 50°C for 10 minutes. Surviving cells were quantified by serial dilution plating (N = 2-3 biological replicates per strain, 142 technical replicates total).

### *R. solanacearum* cells recovered from diseased plant tissue have increased stress tolerance

Bacteria growing inside an infected host can differ significantly in physiology and stress tolerance from cells of the same strain growing in culture(12–14), so we validated decontamination procedures on host-conditioned *R. solanacearum* cells growing in infected tomato plants, a potential source of the pathogen. We heat treated stem sections harvested from symptomatic plants that had been inoculated with each of the seven *R. solanacearum* strains. These stem sections contained around 10^10^ CFU/g tissue as determined by serial dilution plating. Interestingly, about 0.001% of *R. solanacearum* cells inside tomato stems survived 10 min at 50°C; this was true of all seven strains tested. Similar small populations survived stepwise increases in temperature and exposure time. All *R. solanacearum* cells were killed only after infected stems were incubated for 4h at 99°C, with no differences among strains (Fig 5A and data not shown). These results demonstrate that growth in host plants confers dramatically increased heat tolerance on this pathogen.

**Figure 5.**
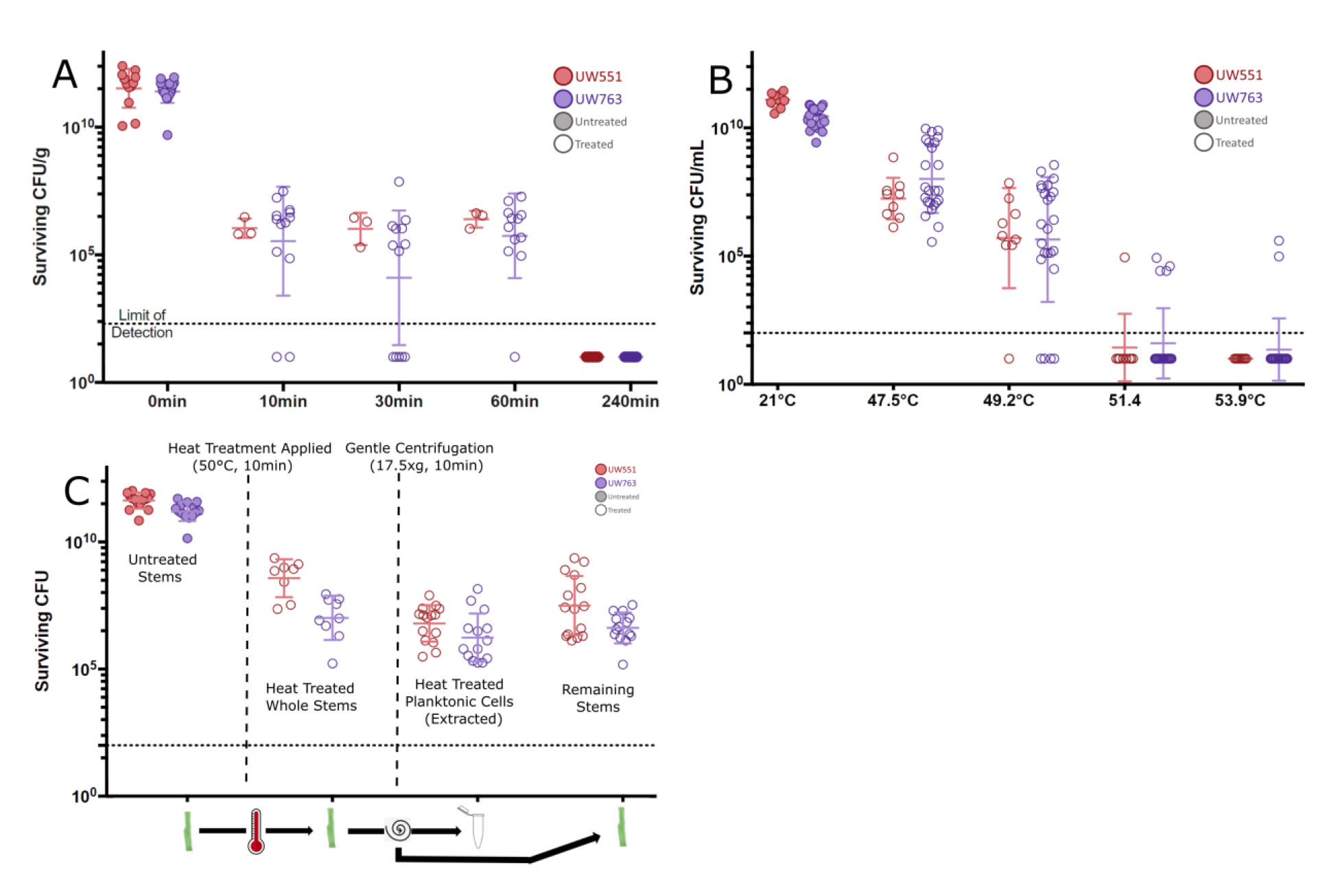
Host-conditioned *R. solanacearum* cells have increased heat tolerance. A 99 °C eradicated *R. solanacearum* cells in plant stems - Population sizes of two *R. solanacearum* strains (indicated by colored symbol) following treatment of infected tomato stems at 99°C for the indicated time. Surviving cells were quantified by dilution plating. (N = 3 biological replicates per strain) **B A small sub-population of host-conditioned planktonic cells have modestly higher heat tolerance** Population sizes of two *R. solanacearum* strains (indicated by colored symbol) following extraction from plant stems and treatment for 10 min at the indicated temperature (N = 3 biological replicates per strain) **C. Host-conditioned planktonic cells heat treated before extraction from plant tissue show increased heat tolerance** Population sizes of two *R. solanacearum* strains (indicated by colored symbol) from left to right: untreated control stems, heat treated whole stem before planktonic cell extraction (CFU/g), extracted planktonic cells (CFU/mL), and remaining stem after heat treatment and planktonic cell extraction (CFU/g) (N = 3 biological replicates per strain with 14-15 technical replicates total).

To further explore how *R. solanacearum* growth *in planta* increased heat stress tolerance we focused on two representative *R. solanacearum* strains, phylotype II R3b2 strain UW551 and phylotype I tomato isolate UW763. Populations of *R. solanacearum* inside wilting plants are heterogenous; some planktonic cells float or swim freely in the flowing xylem sap, while others live in dense biofilms along xylem vessel walls(10, 15). We hypothesized that either: i) bacteria growing in tomato xylem are protected from heat stress by the plant tissue itself; or ii) *R. solanacearum* cells in plants are more heat tolerant because their physiology is altered by the host environment, because their physiology is altered by growth in biofilms, or because they are physically protected by the biofilm matrix.

To test the plant protection and host-altered physiology hypotheses, we measured heat tolerance of planktonic *R. solanacearum* cells following their removal from infected tomato stem sections by gentle centrifugation. Centrifuging *R. solanacearum*-infected tomato stems extracts only about 10% of the total bacterial population(14). If the plant tissue itself offers physical or chemical protection from heat stress, we would not expect extracted planktonic bacteria to survive 10 min at 50°C. However, although most planktonic samples from plants were as heat-sensitive as *R. solanacearum* grown in culture, five out of 33 stems tested contained bacteria with greater heat tolerance, with 15% of replicates across both strains surviving at 51.4 °C, and two of those surviving at 53.9 °C (Fig. 5B). This result could reflect heat-tolerant cells from a biofilm fragment dislodged during centrifugation, or it could indicate that planktonic *R. solanacearum* cells in plants contain a subpopulation of heat-tolerant cells. To further investigate, we first heat treated infected stems for 10 min at 50°C, then centrifuged them to remove planktonic cells and quantified the surviving cells in both the planktonic and stem-associated samples. If the plant offers protection from heat stress, we would expect many planktonic cells to survive treatment. Indeed, we found that some planktonic cells from inside plant tissue survived 10 min exposure to 50°C (Fig. 5C). This supports the hypothesis that the plant offers some protection from heat stress, although it is also possible that heat treatment loosened biofilms inside stems, making it more likely that biofilm fragments were extracted by centrifugation.

To separate the protective effects of growth *in planta* from the protective effects of growing in a biofilm, we measured stress tolerance of *R. solanacearum* biofilms grown on glass slides. If growth in plant tissue protects *R. solanacearum* cells from stress or selects for stress tolerant cells, then biofilms grown on abiotic glass slides would be no more stress tolerant than cells cultured in shaking liquid media. However, post-treatment enrichment culture revealed that cells in biofilms grown on glass slides also survived treatment at 50°C for 10 min (data not shown). This indicated that growth in biofilms on either biotic or abiotic surfaces increased *R. solanacearum* heat tolerance.

Because biofilms can protect microbes from diverse stresses, we hypothesized that *R. solanacearum* living in biofilms in infected plants may also have increased tolerance of UV light, hydrogen peroxide and HuwaSan, but that planktonic cells in xylem sap of diseased plants would have stress tolerance levels similar to those of bacteria grown in broth culture. We tested this hypothesis by subjecting planktonic *R. solanacearum* cells harvested from xylem and resuspended in water at high densities to the physical and chemical antimicrobial treatments described above.

Consistent with our finding that some *R. solanacearum* cells growing *in planta* have increased heat tolerance, *R. solanacearum* cells centrifuged from diseased tomato stems were much more resistant to UV radiation than cultured cells, requiring 5 min exposure to reduce surviving CFU to below the limit of detection, compared to just 30 s for cultured cells (Fig. 6A). These results indicate that *R. solanacearum* cells grown in diseased plants have increased stress tolerance even after the bacteria are physically separated from the plant.

**Figure 6.**
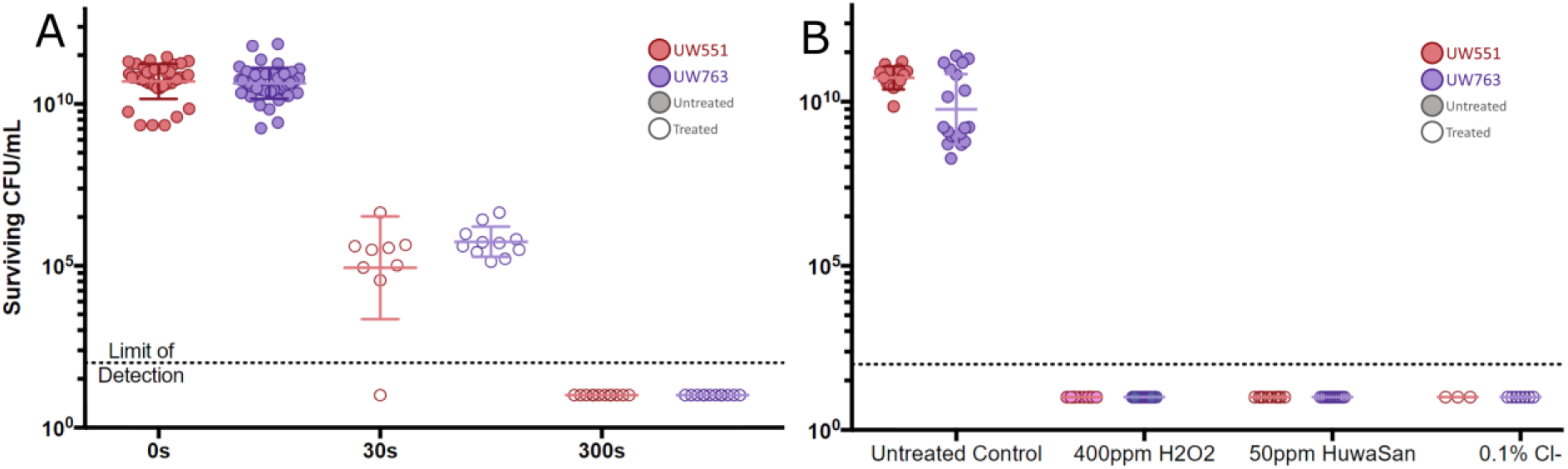
Plant host-conditioned *R. solanacearum* cells have increased UV tolerance, but not increased ROS tolerance. A. Host conditioned planktonic cells show increased UV tolerance - Population sizes of two *R. solanacearum* strains (indicated by colored symbol) following extraction from plant stems and exposure to 0.2 μmol/m^2^/sec UV light on CPG agar plates for the indicated time. Surviving cells were quantified by counting colonies that grew after treatment. **B. Host conditioned planktonic cells do not show increased ROS tolerance** - Population sizes of two *R. solanacearum* strains (indicated by colored symbol) following extraction from plant stems and treatment for 10 min as indicated (N = 3-7 biological replicates per strain) If growth in the plant protects cells from desiccation, this could affect disposal recommendations. To determine if growth *in planta* altered *R. solanacearum* desiccation tolerance, we measured survival of *R. solanacearum* in diseased plant material by thoroughly drying infected tomato stems in a commercial food dehydrator at 35°C. While not strictly equivalent to the 28°C drying that killed all cultured *R. solanacearum* cells, this method ensured uniform and complete desiccation of plant matter which otherwise reabsorbed moisture from air as ambient humidity fluctuated. Two of 121 tested infected stem samples contained detectable surviving cells after 12h at 35°C, a temperature that does not inhibit *R. solanacearum* cells growing in culture. *R. solanacearum* is known to survive for extended periods in soil(5). To determine if exposure to the complex but non-plant soil environment could also increase bacterial desiccation tolerance, we measured *R. solanacearum* survival following complete desiccation of highly infested soil from pots that contained diseased plants. This dried soil contained no detectable *R. solanacearum* cells, suggesting that this pathogen’s tolerance of heat and desiccation is increased by growth in plant tissue, but not by growth in soil.

In response to pathogen infection, plants produce reactive oxygen species (ROS). *R. solanacearum* experiences oxidative stress inside host plants(16). To determine whether growth inside a plant host increased ROS tolerance of *R. solanacearum*, we treated planktonic cells centrifuged from diseased tomato stems with the LD_100_ of bleach, hydrogen peroxide and HuwaSan as determined for cultured cells. Planktonic *R. solanacearum* cells from diseased tomato stems were no more tolerant of these three ROS-generating chemicals than cells grown in culture, indicating that growth *in planta* did not increase *R. solanacearum* tolerance of all stresses (Fig. 6B).

### Inactivation Protocols for Downstream Applications

Diagnostic tests, cloning, and quantifying gene expression all require purified DNA or RNA from R3B2 *R. solanacearum* strains. To comply with Federal regulations that require Select Agents to be inactivated with a validated inactivation protocol prior to downstream applications, we measured *R. solanacearum* survival following four nucleic acid extraction protocols. Representative R3B2 strain UW551 was cultured as described above, then subjected to the cell lysis step of three different commercial nucleic acid extraction kits according to the manufacturer’s instructions or to the lysis step of a phenol-chloroform extraction protocol. Lysate was then serially dilution plated to quantify any surviving cells, followed by enrichment culture to detect any surviving bacteria below the limit of detection. The purified nucleic acid was also spread on rich CPG agar plates and incubated at 28°C for 48 h. No surviving cells were detected in any of these instances, indicating that these four protocols inactivate *R. solanacearum* R3B2.

## Discussion

Our goal was to identify chemical and physical treatments that reliably inactivate *R. solanacearum*, including strains from the highly regulated Select Agent subgroup. Overall, we found that cultured *R. solanacearum* cells are highly sensitive to heat, UV irradiation, oxidative stress, and desiccation (Table 3).

**Table 3:**
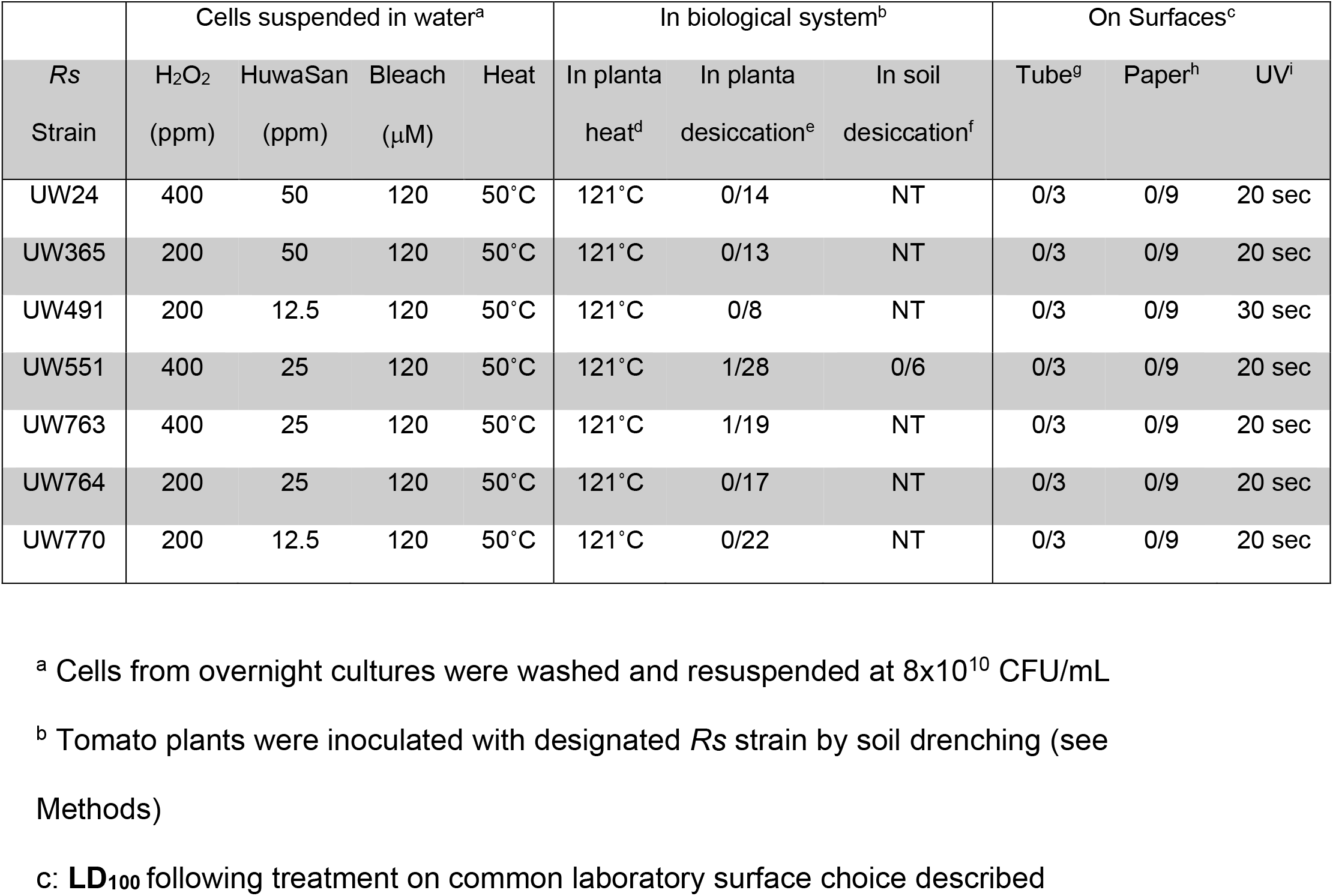
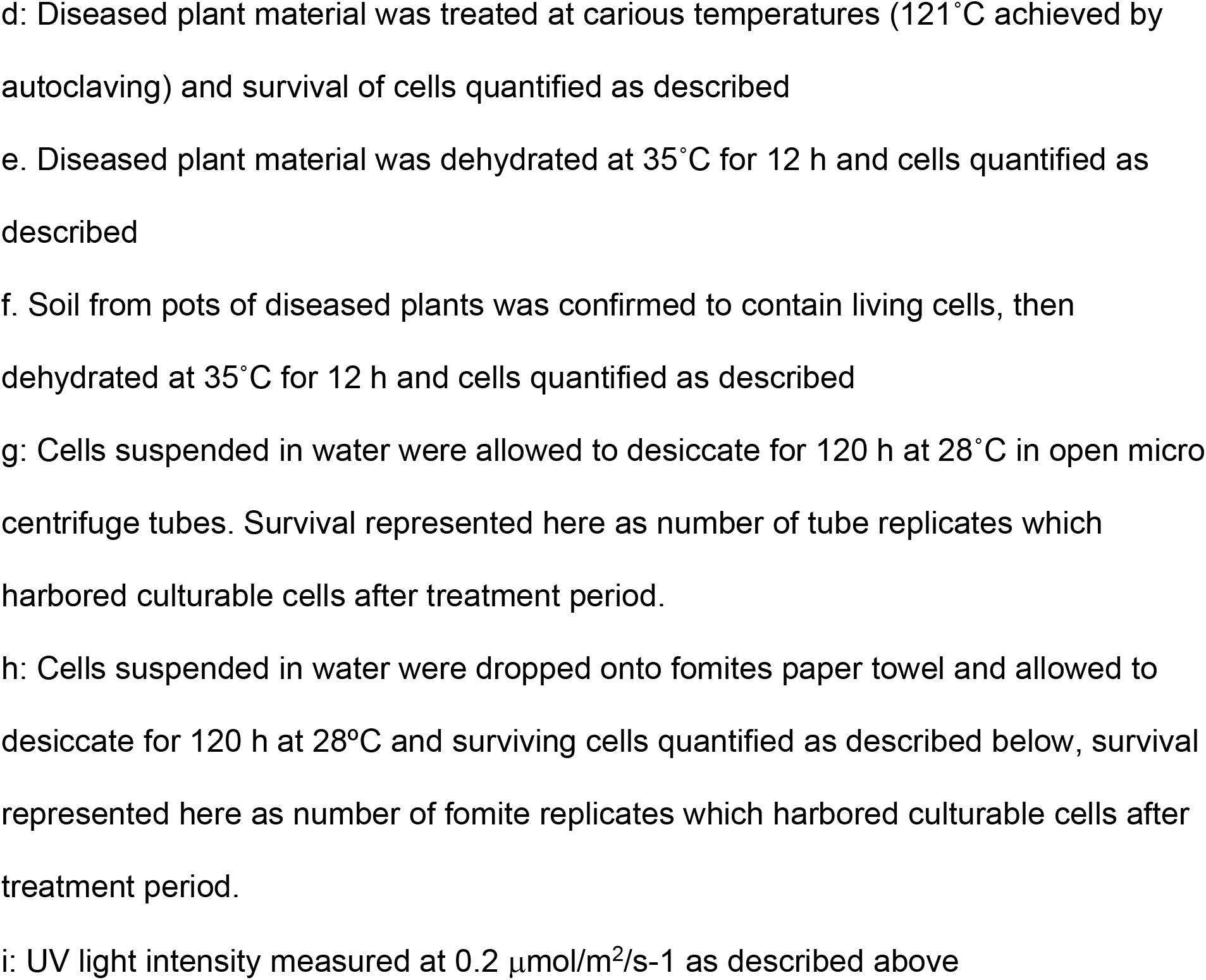
Summary of treatments that eradicated *R. solanacearum* (as LD_100_)

| Rs<br>Strain | Cells suspended in water<sup>a</sup> |  |  |  | In biological system<sup>b</sup> |  |  | On Surfaces<sup>c</sup> |  |  |
| --- | --- | --- | --- | --- | --- | --- | --- | --- | --- | --- |
|  | H<sub>2</sub>O<sub>2</sub><br>(ppm) | HuwaSan<br>(ppm) | Bleach<br>(μM) | Heat | In planta<br>heat<sup>d</sup> | In planta<br>desiccation<sup>e</sup> | In soil<br>desiccation<sup>f</sup> | Tube<sup>g</sup> | Paper<sup>h</sup> | UV<sup>i</sup> |
| UW24 | 400 | 50 | 120 | 50°C | 121°C | 0/14 | NT | 0/3 | 0/9 | 20 sec |
| UW365 | 200 | 50 | 120 | 50°C | 121°C | 0/13 | NT | 0/3 | 0/9 | 20 sec |
| UW491 | 200 | 12.5 | 120 | 50°C | 121°C | 0/8 | NT | 0/3 | 0/9 | 30 sec |
| UW551 | 400 | 25 | 120 | 50°C | 121°C | 1/28 | 0/6 | 0/3 | 0/9 | 20 sec |
| UW763 | 400 | 25 | 120 | 50°C | 121°C | 1/19 | NT | 0/3 | 0/9 | 20 sec |
| UW764 | 200 | 25 | 120 | 50°C | 121°C | 0/17 | NT | 0/3 | 0/9 | 20 sec |
| UW770 | 200 | 12.5 | 120 | 50°C | 121°C | 0/22 | NT | 0/3 | 0/9 | 20 sec |

<sup>a</sup> Cells from overnight cultures were washed and resuspended at 8x10<sup>10</sup> CFU/mL
<sup>b</sup> Tomato plants were inoculated with designated *Rs* strain by soil drenching (see Methods)
<sup>c</sup>: LD<sub>100</sub> following treatment on common laboratory surface choice described
d: Diseased plant material was treated at various temperatures (121°C achieved by autoclaving) and survival of cells quantified as described
e. Diseased plant material was dehydrated at 35°C for 12 h and cells quantified as described
f. Soil from pots of diseased plants was confirmed to contain living cells, then dehydrated at 35°C for 12 h and cells quantified as described
g: Cells suspended in water were allowed to desiccate for 120 h at 28°C in open micro centrifuge tubes. Survival represented here as number of tube replicates which harbored culturable cells after treatment period.
h: Cells suspended in water were dropped onto fomites paper towel and allowed to desiccate for 120 h at 28°C and surviving cells quantified as described below, survival represented here as number of fomite replicates which harbored culturable cells after treatment period.
i: UV light intensity measured at 0.2 $\mu\text{mol}/\text{m}^2/\text{s}$ as described above

*R. solanacearum* survives well in surface waterways, which can be a source of inoculum when they are used for irrigation in agricultural operations(17). Chemical treatments like hydrogen peroxide and bleach are widely used to reduce biological contamination in water(18–20). However, chlorine-containing disinfectants can produce toxic byproducts(21, 22). Hydrogen peroxide is not commonly used in agriculture because it degrades rapidly, can damage infrastructure, and is phytotoxic at high concentrations(23). HuwaSan is an appealing alternative because stabilizing hydrogen peroxide on colloidal silver slows its degradation, making it effective at lower concentrations(24). While our experimentally-determined LD_100_ of 50 ppm differs from the 20 ppm dosage recommended by the manufacturer, we tested much denser bacterial suspensions than those tested previously (10^10^ vs. 10^6^), and we treated for a shorter time (10 min vs. 120 min) (24). Consistent with previous observations, we observed that the LD_100_ of each chemical treatment was affected by bacterial cell density, with lower cell densities requiring lower treatment concentrations. When we tested more environmentally relevant *R. solanacearum* densities of ~10^4^ CFU/mL, all tested dosages above were effective on these smaller bacterial populations (data not shown).

Physical treatments are used in food handling and processing to eliminate pathogens(25–29). Physical treatments can decontaminate while meeting organic standards, which often preclude chemical treatment(30). Though concerns about resistance selection are less common for physical treatments over chemical, the mechanism of action of physical treatment is often not fully understood. We tested the efficacy of three physical treatments: heat, desiccation, and UV irradiation.

We found that *R. solanacearum* cultured in rich media was completely intolerant of even brief (10 min) exposure to temperatures over 50°C, meaning that heat treatment is a reliable way to eradicate laboratory-grown cells of this pathogen. However, growers and regulators must dispose of plant material, not cultured bacteria. Unfortunately, growth in a plant host dramatically increased heat tolerance of *R. solanacearum* cells, which survived for at least 18 days inside diseased plant material at 50°C (data not shown).

Diseased plants represent a release risk and agricultural operations rarely have the infrastructure to autoclave or incinerate large amounts of plant material. It is unclear whether *R. solanacearum* cells liberated from plant material via decomposition would maintain an increased heat tolerance, or how heat tolerance interacts with competitive fitness in a microbial community. Measuring the survival of the pathogen in an agriculturally relevant environment such as hot compost piles may answer this question.

In host plant xylem vessels *R. solanacearum* forms biofilms. These robust biological matrices are linked to the pathogen’s fitness and virulence (Khokhani et. al. 2017). If the increased heat tolerance of *R. solanacearum* growing *in planta* is due to protection by the biofilm or some other physical or chemical condition in the plant itself, cultured cells and planktonic (non-biofilm) cells from plants should be similarly sensitive to heat. We observed limited survival of planktonic cells extracted from plants before treatment at 51° or 54°C. However, planktonic cells that were first heat treated inside plant stems, then extracted and evaluated for survival had much higher survival rates. This supports the hypothesis that being inside a tomato stem makes *R. solanacearum* more heat tolerant. To probe the question of whether the bacterium must be host-conditioned to be more stress tolerant, we also measured survival of heat-treated *R. solanacearum* cells in *ex vivo* biofilms on glass slides. If bacteria in biofilms are protected from stress, biofilms grown *ex planta* should also offer protection. Indeed, *R. solanacearum* cells in biofilms on glass slides did survive 50°C heat treatment for 10 min. This supports the hypothesis that *R. solanacearum* cells in biofilms are physically or physiologically protected from stress. This is consistent with observations that growth in biofilms increases stress tolerance of many bacteria(31).

High-energy ultraviolet (UV) light is commonly used in clinical settings to eliminate pathogen transmission on surfaces(32). This approach is also used to decontaminate agricultural equipment and the surfaces of plants during growth and foods after harvest(33–35). UV light is also a common water decontamination method, for application at both small scales (personal water filters) and large scales (irrigation water treatment)(36). Because *R. solanacearum* can be transmitted and dispersed through contaminated equipment, we tested eradication efficacy of UV light by treating cells on agar plates. This proved to be highly effective. However, we did observe that any cells that were protected from exposure (e.g. stabbed into the agar surface with a pipette tip) survived irradiation. This suggests direct exposure to UV light is necessary to decontaminate equipment. Further, host-conditioned *R. solanacearum* from stems of infected tomato plants were much more tolerant of UV radiation. Similar ‘persister cells’ have been observed in clinical pathogens under antibiotic pressure(12), and represent an avenue of future study. Unfortunately, implementation of physical treatment methods on a large scale requires costly infrastructure modifications such as in-line UV treatment of irrigation water (discussed below). However, the advent of small, portable, and highly efficient LED and visible spectrum irradiation equipment may make this eradication method more accessible to small operations(35, 37).

To demonstrate practical utility of these eradication methods, we tested three *R. solanacearum* phylotype I strains isolated from sites currently experiencing bacterial wilt outbreaks. We field-tested effectiveness of some methods against strain UW763, which came from a large export tomato operation in Senegal that was suffering serious wilt disease losses. The strain was isolated from symptomatic plants and from the farm’s irrigation water source. Treating irrigation water with 25 ppm HuwaSan and an in-line UV irradiation system drastically reduced bacterial wilt disease incidence in this farm (confidential personal communication). Further, using centrifugal concentration followed by enrichment culture we detected no *R. solanacearum* cells in treated water samples from this facility during regular monitoring over 18 months. This indicates that the treatments described here can be effective in field conditions.

We also showed that *R. solanacearum* cells are killed by the lysis step of four different nucleic acid extraction protocols. These validated inactivation protocols satisfy the CDC-USDA requirement that Select Agent research labs experimentally verify that whole agent or subcellular components are fully inactivated (killed) before being removed from registered Select Agent research lab space(7). This facilitates downstream biochemical applications such as DNA-based diagnostic tests or analysis of proteins or gene expression. The cell lysis step of other extraction protocols would probably also inactivate *R. solanacearum*, but this would need to be experimentally verified using the procedures described here.

We observed minor or no differences among our set of seven *R. solanacearum* strains with respect to LD100 for any tested eradication method (Table 3). To explore possible variations in susceptibility among the highly regulated R3bv2 subgroup, we quantified survival of four different R3bv2 strains. There was no evidence that members of the R3bv2 subgroup were more difficult to eradicate than other *R. solanacearum* strains. This means the same practices can be safely used on all *R. solanacearum* strains, regardless of whether the sub-classification of the infesting strain is known.

Practical and scalable methods to control *R. solanacearum* are needed to increase security of high value export and subsistence crops. The eradication protocols validated here can be used by growers to control infestations and reduce crop losses to bacterial wilt. They will also support regulators charged with developing phytosanitation and biosafety protocols, and researchers who must comply with regulations and prevent accidental release of *R. solanacearum*.

